# Understanding behavior in a captive Lion-tailed macaques (*Macaca silenus*) group

**DOI:** 10.1101/692723

**Authors:** Jayashree Mazumder

## Abstract

A captive lion-tailed macaque group, consisting of two adult males and one adult female, was observed over a period of three months. We collected the data from 8.30hrs to 17.30hrs, using both focal and scan animal sampling methods. In the study, we divided the behaviors into abnormal and normal behavioral traits which were further divided into self-directed and social interchange behaviors. When compared with the past records on the wild and captive group, most of the behaviors were similar which included behaviors like forage, grooming, aggressive display and reproductive behavior. Animals who were born in a zoo or confiscated from another zoo showed higher levels of abnormal behaviors. The results of this study discuss the range of behavioral patterns displayed by the captive animals, discuss the causal factors for the behavioral pathogens, and further suggests feasible solutions to increase the welfare of these macaques. In the study, the early life history of the animal and the lack of social and environmental stimulus could be very critical for the development of behavioral pathogens. However, to establish this would require more research.

## Introduction

Lion-tailed macaques (*Macaca silenus*) are endangered species (Kumar, Singh, & Molur, 2008) and they are endemic to the Western Ghats in Southern India (Mallapur, Sinha, & Waran, 2005). This species lives in a group of 8 to 49 individual with more number of females and only 1-3 males i.e. multiple male-multiple females (Kumar, 1987; Raghavan, 2001). They are mostly frugivorous, but they are also found to consume insects and birds (Mallapur, Waran, & Sinha, 2005a). They are mostly arboreal in nature and live in the rainforests of the Western Ghats, India (Kumar et al., 2008). The forest of southern India has a complex structure with trees forming canopies and water bodies passing through the forests (Mallapur, Sinha, et al., 2005; Mallapur, Waran, & Sinha, 2005b). To house a species living in such ecologically rich environment, the captive environment must also be equally enriched. Nevertheless, the condition of the Indian zoological parks housing lion-tailed macaques seems to be barren with no ecological or social stimulating environment, which restricts the animal biologically from displaying species-specific forms of behavior (Mallapur, Waran, et al., 2005b). Captive environment enforces an artificial environment on the animal which differs greatly from their natural habitat. The amount of behavioral changes the species undergo while living in captivity depends on the enrichment provided in the enclosure (Mallapur & Choudhury, 2003).

Stress and boredom are two important phenomenon inducing behavioral pathogens in captive animals (Mallapur & Choudhury, 2003). Lack of social interaction results in the development of behavioral pathogens (Kaumanns, Singh, & Sliwa, 2013; Mallapur & Choudhury, 2003; Raghavan, 2001). Behavioral abnormalities have been seen in the form of self-mutilating behaviors, aberrant sexual activity and stereotypic activities which are some of the most common behavioral pathologies among animals living in a captive environment with no social bonding (Mallapur & Choudhury, 2003; Mallapur, Waran, et al., 2005a). The inability of the individuals living in the zoo to adapt to their captive environment severely compromises the welfare and health of the individuals (Mallapur, Waran, et al., 2005a). In the contemporary world, a lot of focus has been diverted outside India towards conducting research on the primates found in captive environment to study the factors influencing their behavior, health and reproduction to formulate proper measurements and welfare (Mallapur, Waran, et al., 2005a; Mitchell et al., 1988; Mitchell, Soteriou, Towers, Kenney, & Schumer, 1987). Social deprivation is on the key factors which affect the behavior of the captive individuals, particularly nonhuman primates (Hosey, 2005; Khan, 2013; King, Stevens, & Mellen, 1980; Mallapur & Choudhury, 2003). Highest degree and frequency of abnormal behavior and poor welfare have been documented among captive individuals with a rearing history in isolation (Anderson & Chamove, 1980; Mallapur & Choudhury, 2003; Mallapur, Waran, et al., 2005a; Mootnick & Baker, 1994). Self-mutilation has been observed among single-housed animals and among animals who have been deprived of social stimuli (Mallapur, Waran, et al., 2005a). Such traits are restricted to zoo environment and are thus very rarely or almost never found among free-ranging animals or animals living in proper social groups (Anderson & Chamove, 1980; Chamove, Anderson, & Nash, 1984; Mallapur & Choudhury, 2003; Mallapur, Waran, et al., 2005a). The enclosure design also plays a very important role in influencing the behavior as primates requires large area of usable space both horizontally and vertically which is complex in nature providing all the necessary stimuli which would kindle them to exhibit their species-specific forms of behavioral patterns (A. S. Clarke & Lindburg, 1993; A. Susan Clarke, Juno, & Maple, 1982; Goerke, Fleming, & Creel, 1987; O’Neill, Novak, & Suomi, 1991). A distinguishing behavioral difference among free-ranging and captive individuals is that free-ranging animals spend most of their time foraging and gathering food travelling a long distance within their territory, but the zoo primates are devoid of such opportunities as they are provided food at definite durations and thus they do not require any kind of foraging or gathering activity (Mallapur, Waran, et al., 2005b).

In the absence of the environmental and social stimuli, unusual behavioral forms are developed which are not natural (Bloomsmith, Alford, & Maple, 1988; Mallapur, 2005a; Marriner & Drickamer, 1994). In this article, we aim to carefully examine the range of social behavior and discuss the abnormal behaviors displayed by captive lion-tailed macaques in National Zoological Park (NZP), New Delhi, India. Further, we discussed the causal factors and suggested methods for reducing the stress among these individuals and other such animals who exhibit behavioral abnormalities.

## Method

### Subject

This behavioral study was conducted among three captive lion-tailed macaques housed in National Zoological Park (NZP), New Delhi. The behavioral observation was done for a duration of three months from December 2015 to February 2016, in addition to a pilot survey done in the month of October and November 2015. The NZP, New Delhi house five lion-tailed macaque species, but only three were present in the enclosure during the study period. Since the mother with her newly born infant was kept in the veterinary hospital and research on them was not permitted, the present study was conducted on two adult males and one adult female present in the enclosure. One among the two males was designated as the alpha male (named as P.K.) who was morphologically the largest in size and had an abnormal fat deposit in his abdominal region. The other male was comparatively thin and was named as Arun. The female was the smallest in size and was named as Mrinal. According to the zoo records, P.K. was wild-caught, whereas Arun was confiscated and Mrinal was zoo-born (Fig. 1).

**Fig. 1:**
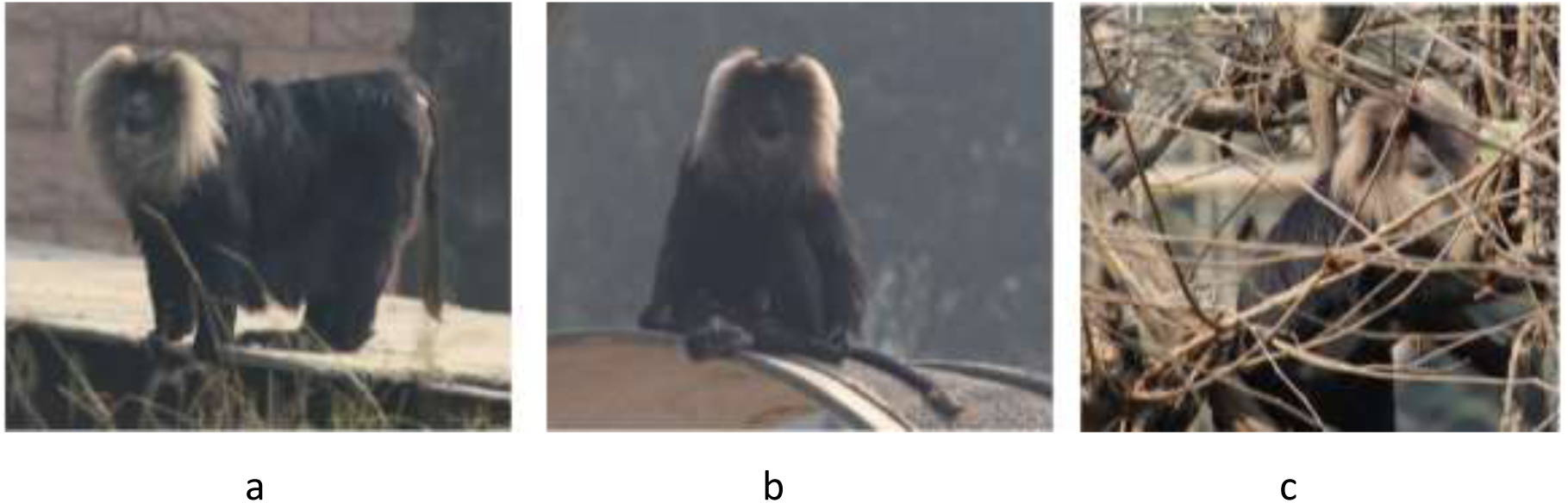
The animals studied (a) P.K., (b) Arun (c) Mrinal.

### Enclosure

The lion-tailed macaques exhibit was large with an approximate radius of 5m and was surrounded by a 3m wide water body. The entire enclosure was surrounded by a water dyke and is a typical wet moat type of enclosure (Fig. 2). The enclosure had a soft ground and was semi enriched as it has few poles with some ropes, but the trees planted was nothing like the canopy-forming trees in the rainforest and barely had leaves on their branches. Metal platforms and hollow post were put up within the enclosure. The entire exhibit was divided into visitor area (for visitors only), edge zone (the side near to the animal shelter, with minimal accessibility to view animals), off-exhibit area (covered area with no accessibility to view animals) and the back zone (entry for the zoo care-takers to the animal shelter where the animals were kept after the visiting hours). The area was enriched by establishing trees, poles, sleeping shelters and an arch. The enriched zone was further divided into enrich zone (area with more variability of enrichment) and other zone (area with only poles connected with each other by ropes). The enclosure was connected to the cage by a ramp or a bridge. The entire enriched zone was accessible to the animals, but the movement of the animals was restricted to enrich zone except for Arun who was more visible in the other zone and mostly foraged through the banks of the water dyke. Mrinal was seen mostly near the ramp, whereas P.K. spent most of his day sun-bathing and sleeping on the arch or the shelter. The shelter was solely used by P.K. whereas the arch was rarely seen to be used by him. Mrinal spent most of his day hiding under the ramp and was also found clinging to the cage door.

**Fig. 2:**
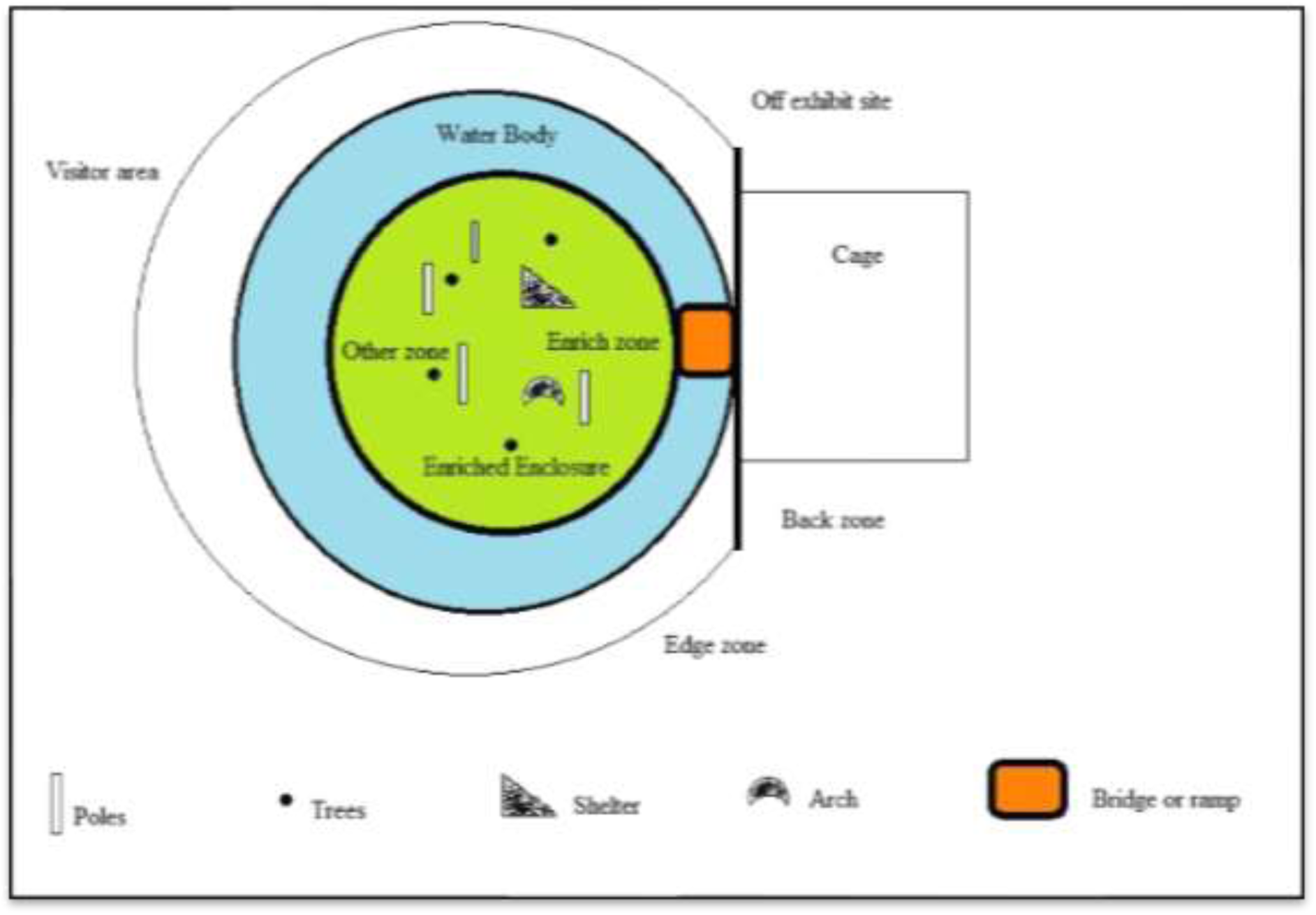
The enclosure zones marked on a base map for the outdoor lion-tailed macaque

**Fig. 3:**
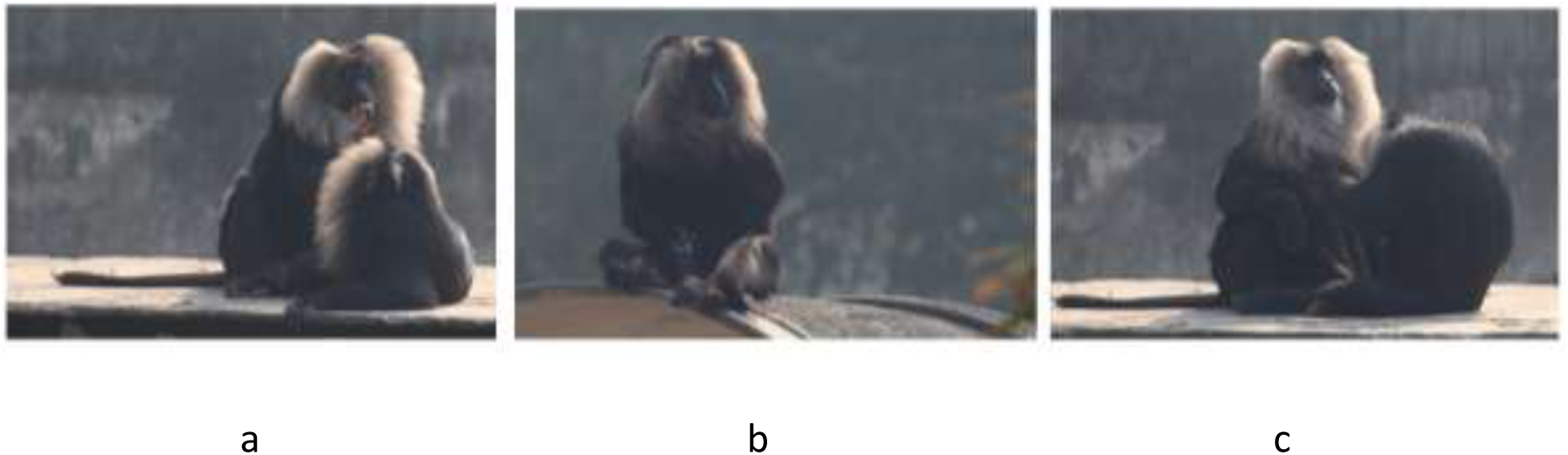
The individual behaviors observed during the study (a) P.K. threats Arun, (b) Arun masturbating, (c) Mrinal grooms P.K

### Procedure

During the pilot survey *ad libitum* sampling was carried out to record the types of behavior displayed by the members in the group (Altmann, 1974). In order to standardize the recording procedure of the behaviors, ethogram has been prepared. The description of the ethogram has been done in the following section. In the later stage focal and scan sampling was conducted to obtain qualitative and qualitative data and, on an average, every individual was studied for 10.6 ± 5.4 hours/ individual for a total duration of 26 hours over a period of 3 months. The study was conducted throughout the day starting from 08.30h in the morning till the zoo closed around 17.30h, throughout the week excluding Fridays since Fridays NZP remained closed for public. Each sampling was instigated with scan sampling of all the members in the enclosure followed by a focal animal sampling of a predetermined individual within a group for a duration of 15 minutes (Mallapur, Waran, et al., 2005a). Every subsequent focal animal sampling was followed by scan sampling and then a focal sampling of the other individual within the enclosure. The individuals were sampled in an orderly manner and it was decided based on the observations done on a daily basis prior starting the sampling sessions. Names were allotted to each individual in the group. Using random sampling the order in which the members will be studied was chosen. This was done by writing the names on pieces of paper, which were then put in a box and shuffled. The names were then randomly picked out thereby proving each member a position in the queue for focal animal sampling. For every day, a new order was obtained in the same manner. The entire set of behavioral events have been expressed in terms of number of events per hour spent by an individual. The data for all the three individuals were pooled at the end of the observation period to obtain the group averages.

## Results

The captive animals displayed 27 normal behavioral traits which are very commonly seen among the wild animals (Graph 1). When compared out of the 27 behavioral variants, P.K. displayed 23 normal behavioral traits (85.1%), followed by Arun who displayed 21 behavioral traits (77.7%) and Mrinal had showed least abnormal traits 16 (59.2%) (Graph 1).

**Graph 1:**
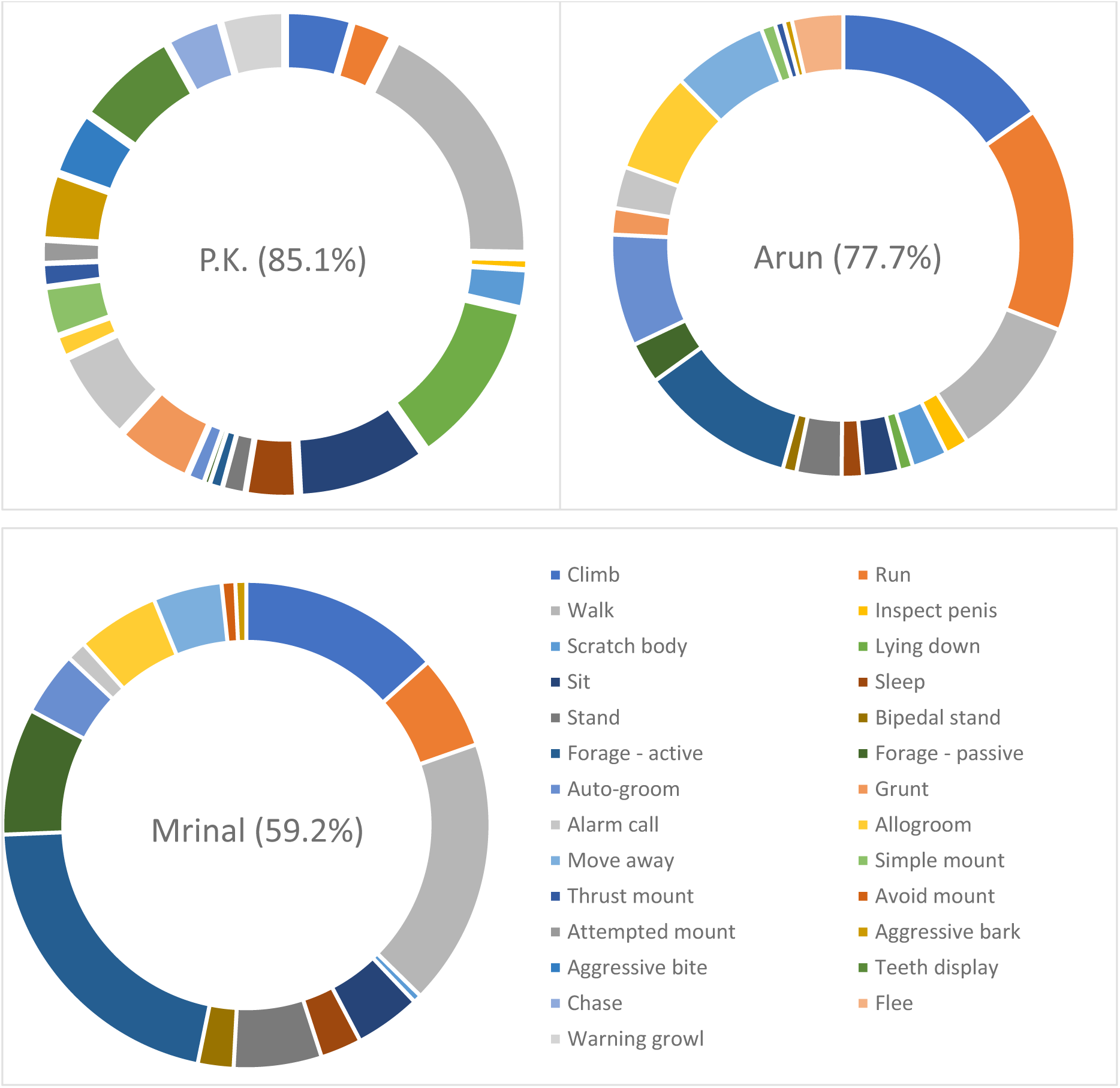
Normal behaviors displayed by the lion-tailed macaques at NZP

The captive lion-tailed macaques showed 40 behavioral traits and individually P.K. showed 26 patterns, Arun displayed 32 patterns whereas Mrinal was observed to display 22 behavioral patterns (Table 1).

**Table 1:**
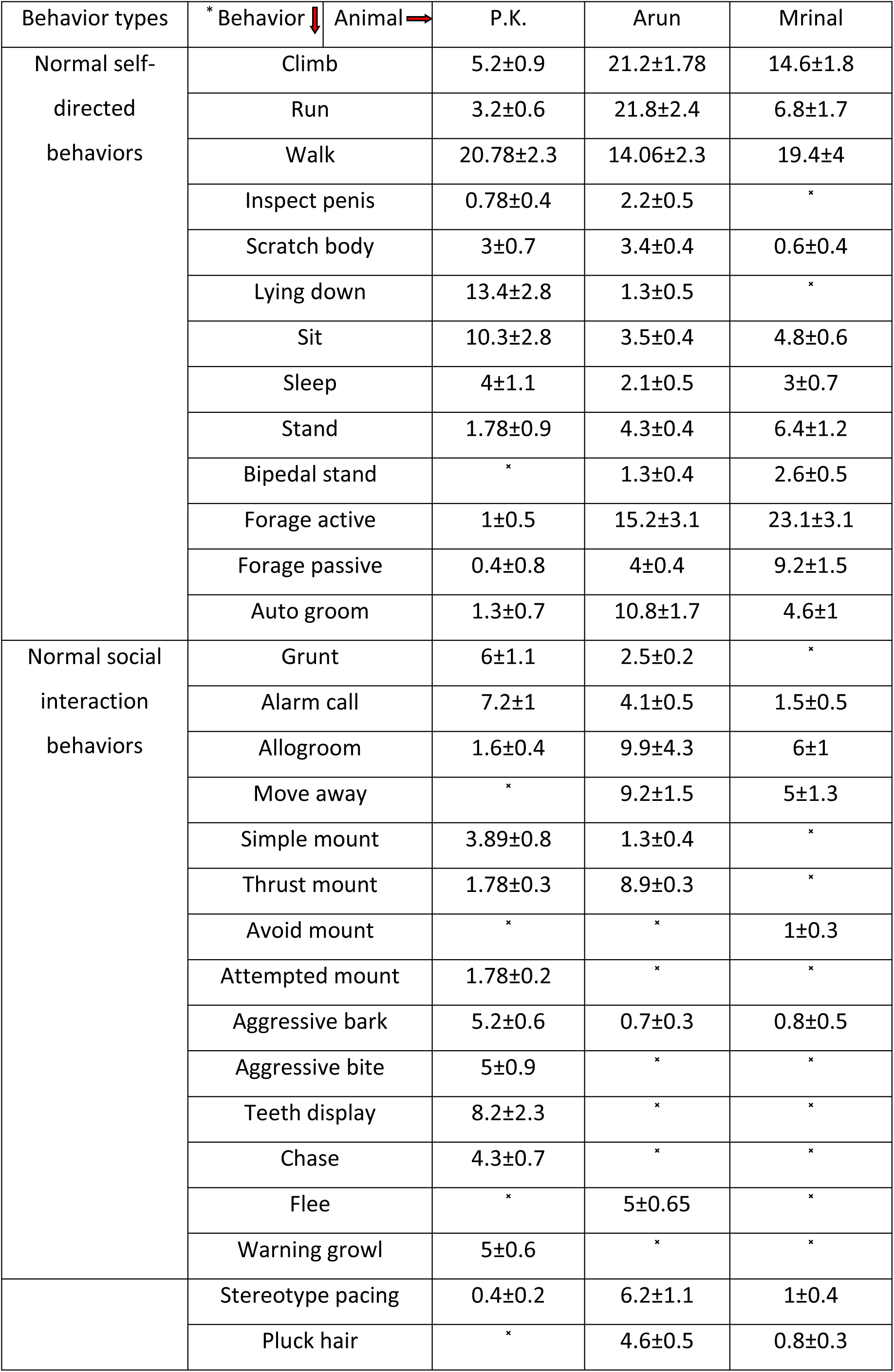
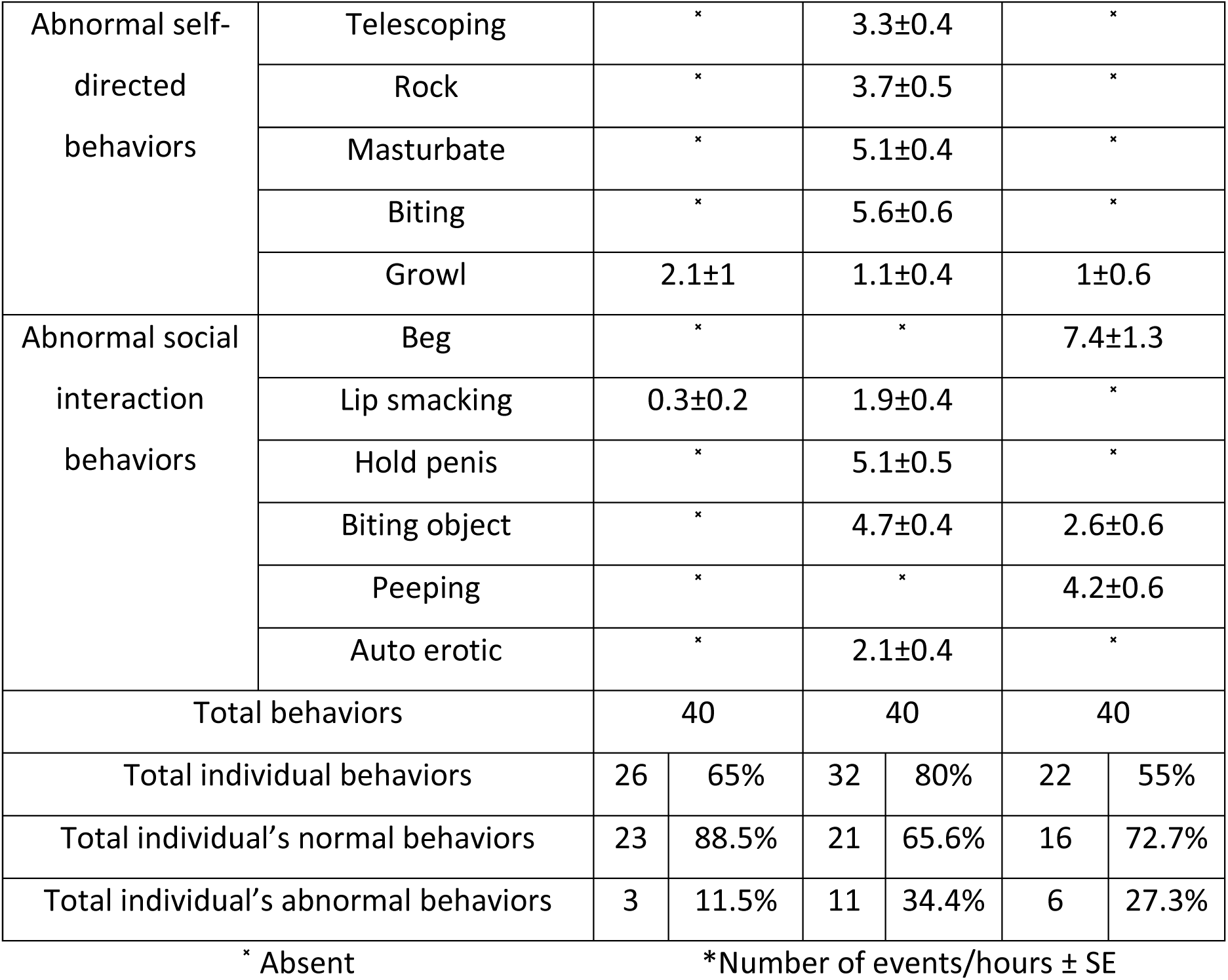
Distribution of behaviors among the captive individuals at NZP (^×^ Absent)

The captive lion-tailed macaques displayed 13 abnormal behavioral traits (Graph 2), which could be divided into seven self-directed (Graph 5) and six-social interchange (Graph 6). When the behaviors are compared between the individuals, Arun displayed the highest variety of behavioral patterns. Out of 13 abnormal behavioral traits, 11-behavioral traits (84.6%) was displayed by Arun alone whereas Mrinal showed six behavioral traits (38.4%) and P.K. showed the least variability in terms of abnormal behavior display i.e. three-behavioral traits (23%). Lip smacking and growl was visible in all the three individuals with P.K. showing the highest rate of growling behavior (Graph 2).

**Graph 2:**
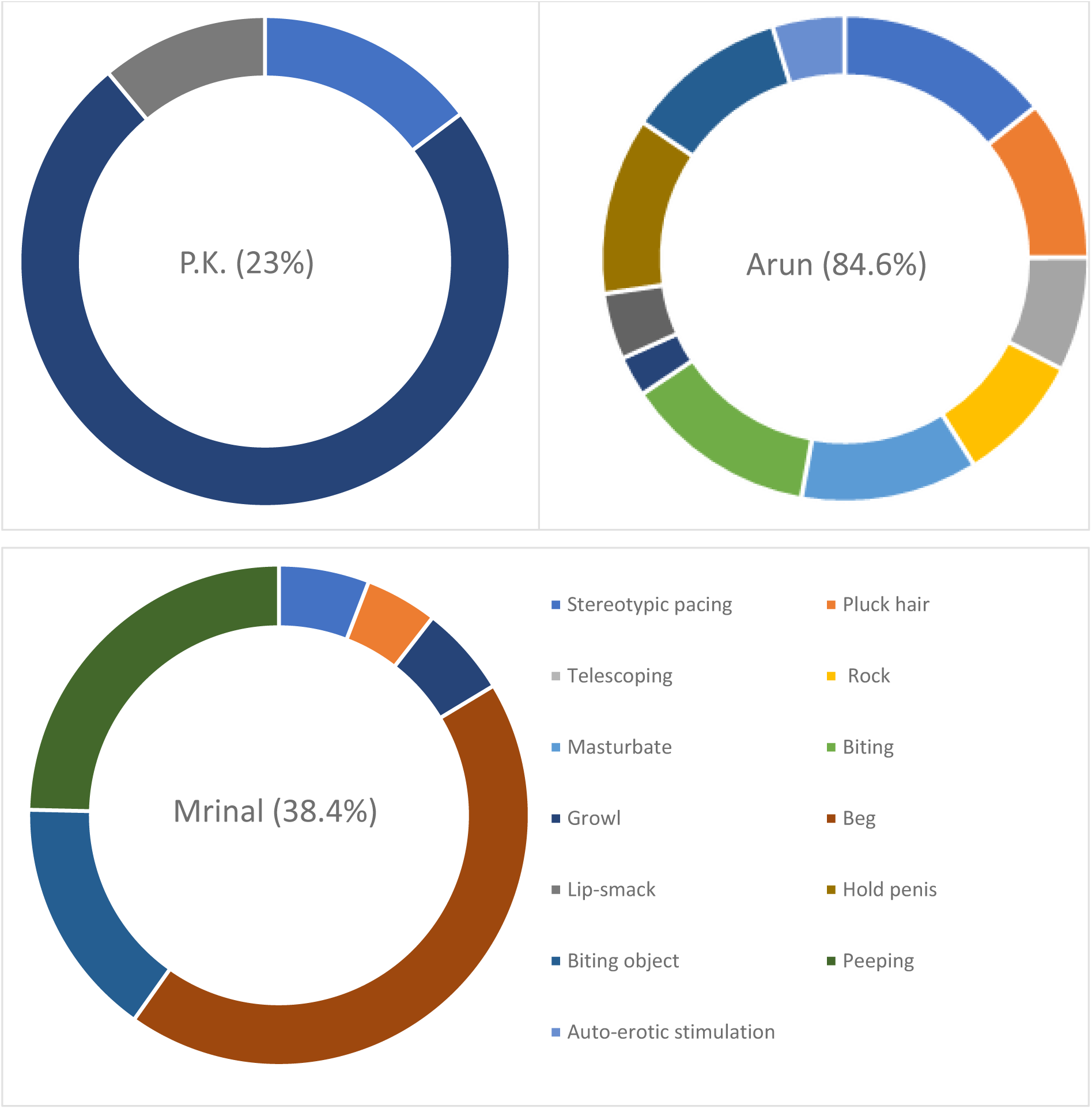
Abnormal behaviors displayed by captive lion-tailed macaques at NZP

When the behaviors are compared, P.K. exhibited a higher level of normal behavioral pattern than Arun. Some of the behaviors were unambiguously done by particular individuals only. Behaviors like beg and peeping were observed only in the female, whereas Arun was observed to display a larger variety of abnormal behaviors as telescoping, rock, masturbate, biting, holding penis and autoerotic behavior (Table 1).

Overall, Arun showed high frequencies of behavioral variants, followed by P.K. and then by Mrinal (Table 1). And when the behaviors are compared, Arun also showed high rates of abnormality than the other two members (Table 1). When compared between individuals for normal and abnormal behavioral distribution, P.K. shows comparatively less abnormal traits than others (Table 1).

A proper analysis of each behavior shows that P.K. spent most of the time displaying resting behavior, whereas Arun and Mrinal were actively indulged in foraging activities. Arun spent most his time by grooming himself or by inspecting his penis, he was very rarely seen to be lying down during the observation period (Graph 3). When comparing the data of normal social interaction behaviors, P.K. was found to be very active during social interactions between the group members. Alarm calls were mostly initiated by P.K. and he actively indulged in territorial protection and displayed aggression in the form of bark and teeth display (Graph 4). On one occasion, a langur was seen to encroach on the enclosure, P.K. actively participated in giving threat calls, and Mrinal followed him behind. Arun on that occasion was seen hiding behind Mrinal with very minimal alarm calls.

**Graph 3:**
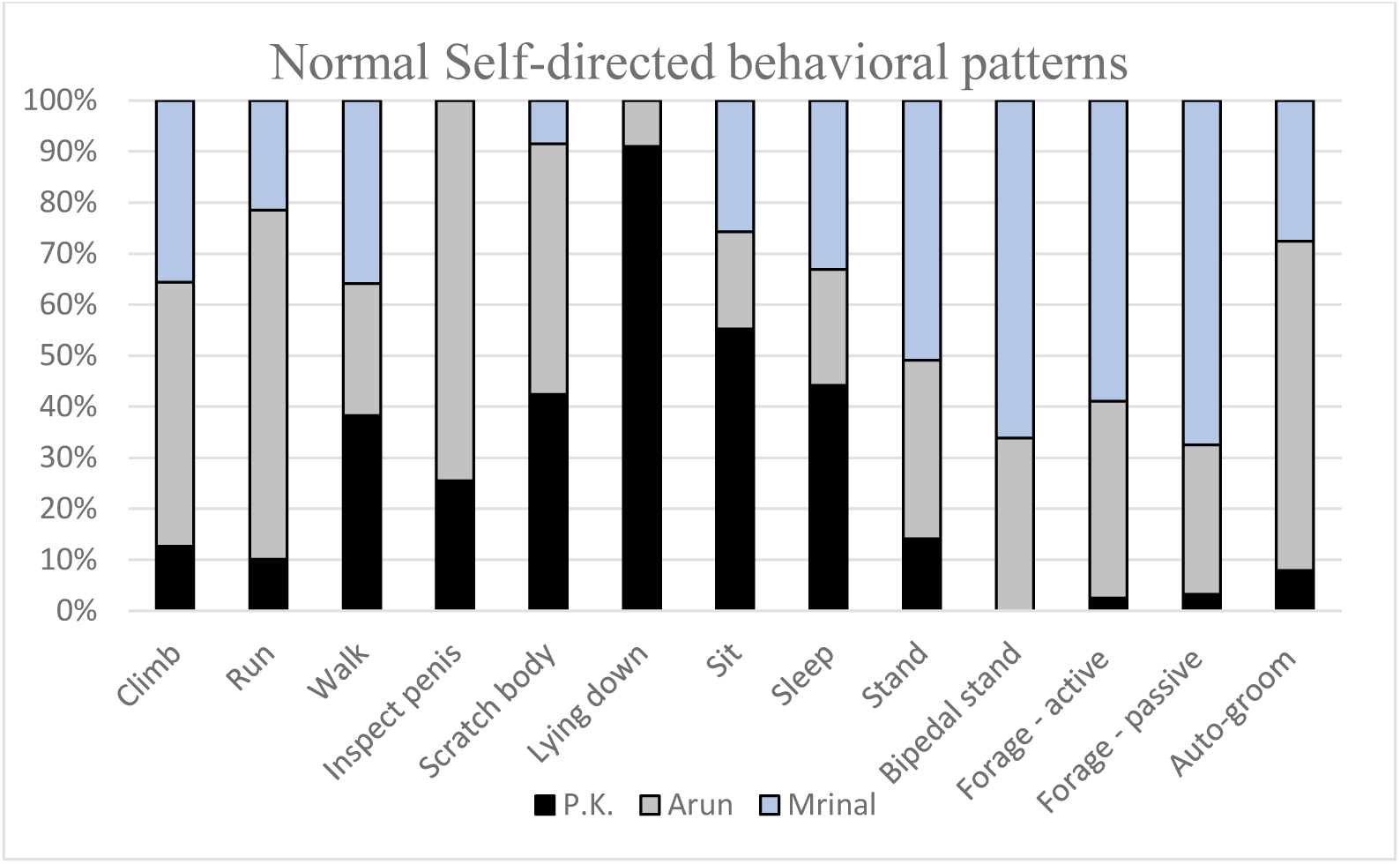
Normal self-directed behaviors by the lion-tailed macaques at NZP

**Graph 4:**
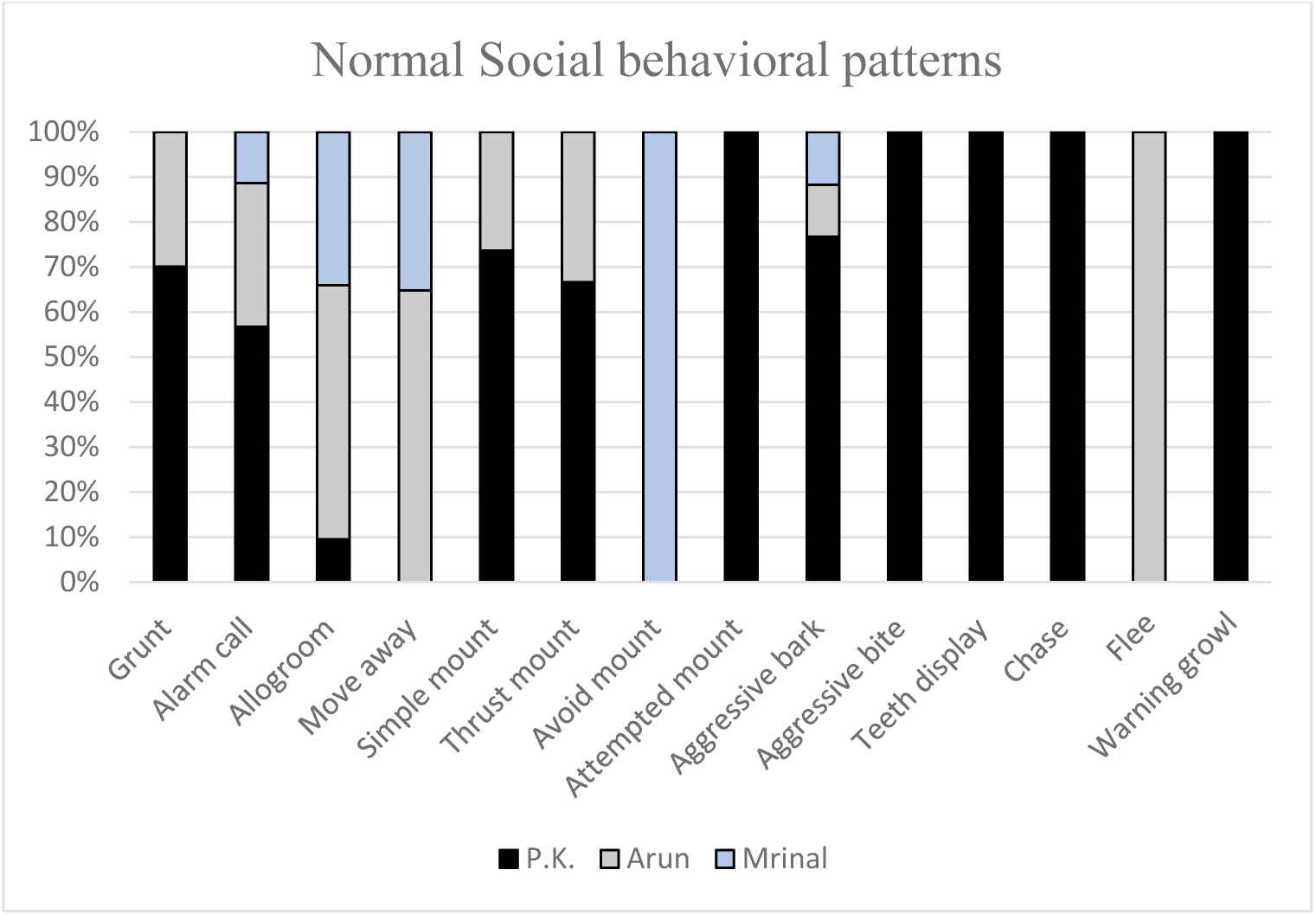
Normal social behaviors by the lion-tailed macaques at NZP

Arun was the only individual observed to display behavior like telescoping, rock, masturbation and biting (self-biting) behavior. Stereotype pacing and growl were observed in all the three individuals, but higher rates of stereotype pacing were observed in Arun, whereas growl was observed in P.K (Graph 5).

**Graph 5:**
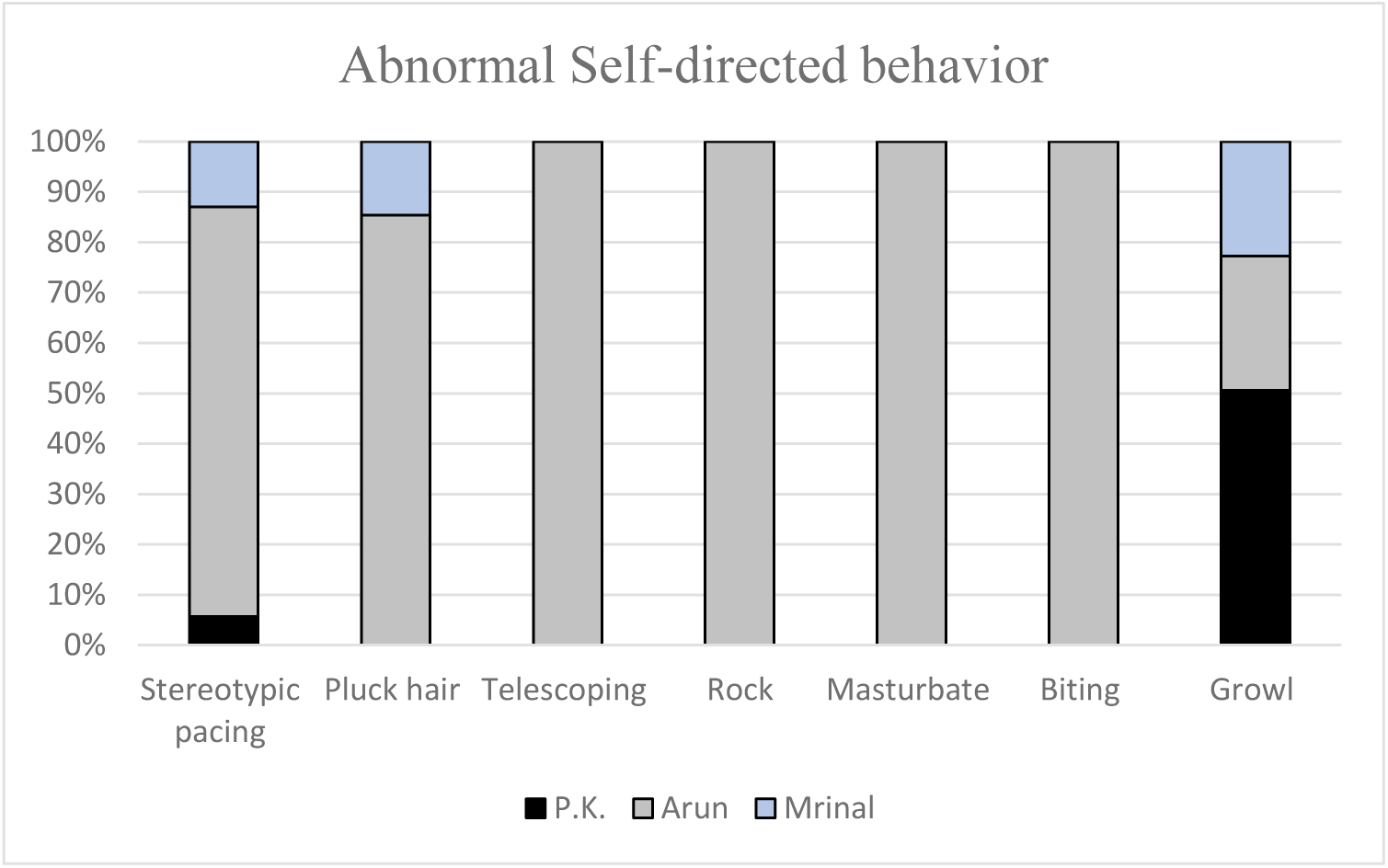
Abnormal self-directed behaviors by the lion-tailed macaques at NZP

Social interaction data depict behavioral pathogens among Mrinal and Arun. Beg and peeping were observed in Mrinal alone, whereas behavioral variants as holding penis and auto-erotic stimulation were observed by Arun. P.K. and Arun were observed to display lip-smacking behavior (Graph 6).

**Graph 6:**
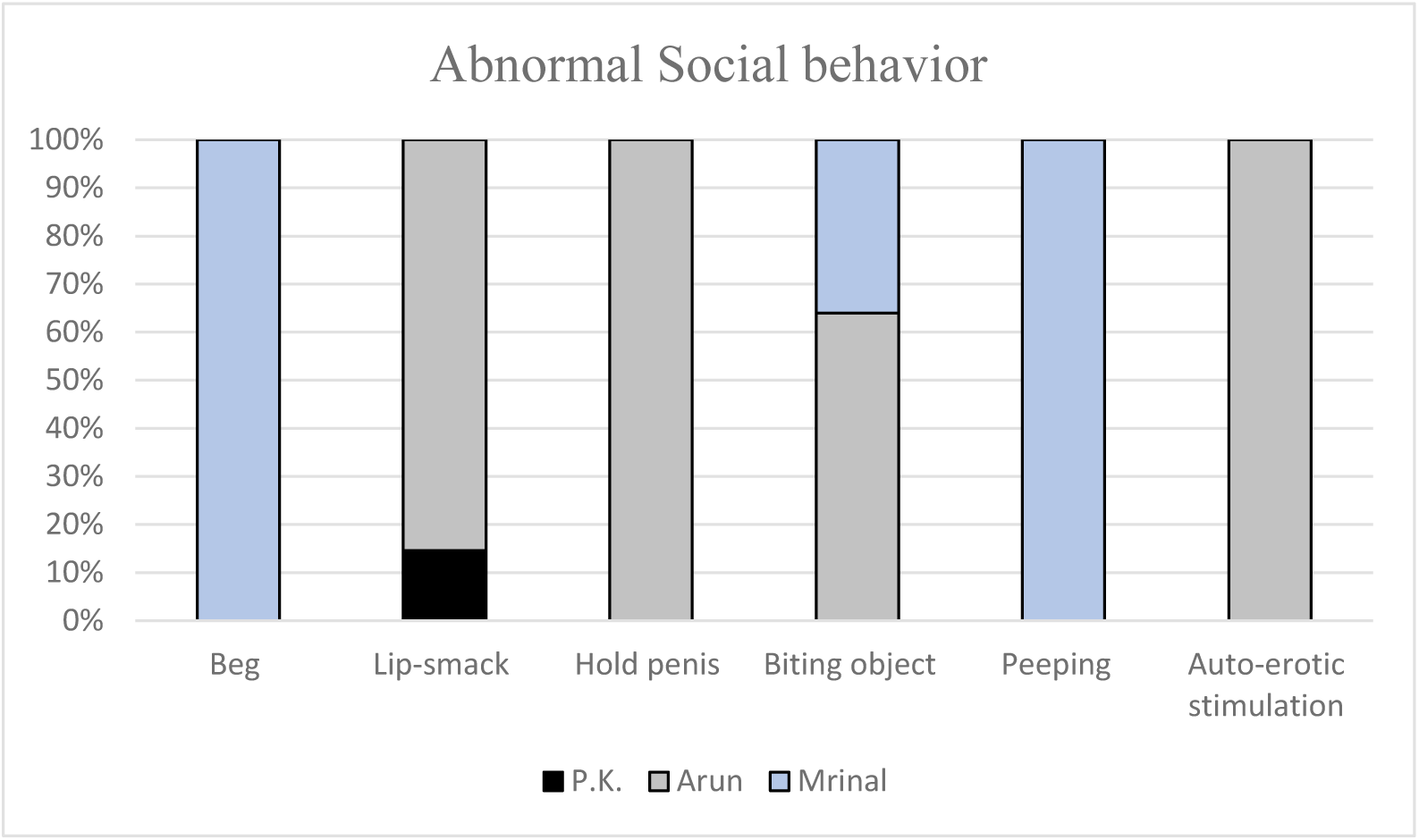
Abnormal social interaction behaviors by the lion-tailed macaques at NZP

Comparing the social behavior data among the individuals, aggressive behavior (threat, lip smacking) was more between Arun and P.K. The reproductive behavior (mounting behavior) was displayed by P.K. towards Arun as a nature of depicting dominance hierarchy and towards Mrinal as a sexual preference. P.K. was never seen to avoid any kind of mounting behavior. Arun also attempted mounting behavior but was avoided by Mrinal. Social behavior like grooming differed between the individuals. Mrinal groomed P.K. and was also groomed by him. Arun was seldom groomed by Mrinal but he actively groomed P.K. and occasionally groomed Mrinal. During alarm calls, calls initiated by P.K. was given highest preference and all members followed behind, except for sometimes when Arun stayed silent. Alarm call initiated by Arun was mostly replied by P.K. and Mrinal avoided any type of social contact with him (Graph 7). This avoidance of Mrinal towards Arun and occasional threat from P.K. resulted in social isolation, which increases Arun’s stress. Thus, lacking social bonding made him innovate and he was observed to open screws and dismantling a hollow pole by trial and error process. He would spend most of his time foraging and during the later morning half, he would either try to break open a pole or try to get hold of a screw on top of the log. Although, we could not decide the exact reason for Arun to get indulged in such activities, lack of social interactions is widely seen to be the reason behind the development of such behaviors, because such behavior was not seen in the other two individuals who would groomed each other or display resting behavior while Arun engaged in disassembling the poles.

**Graph 7:**
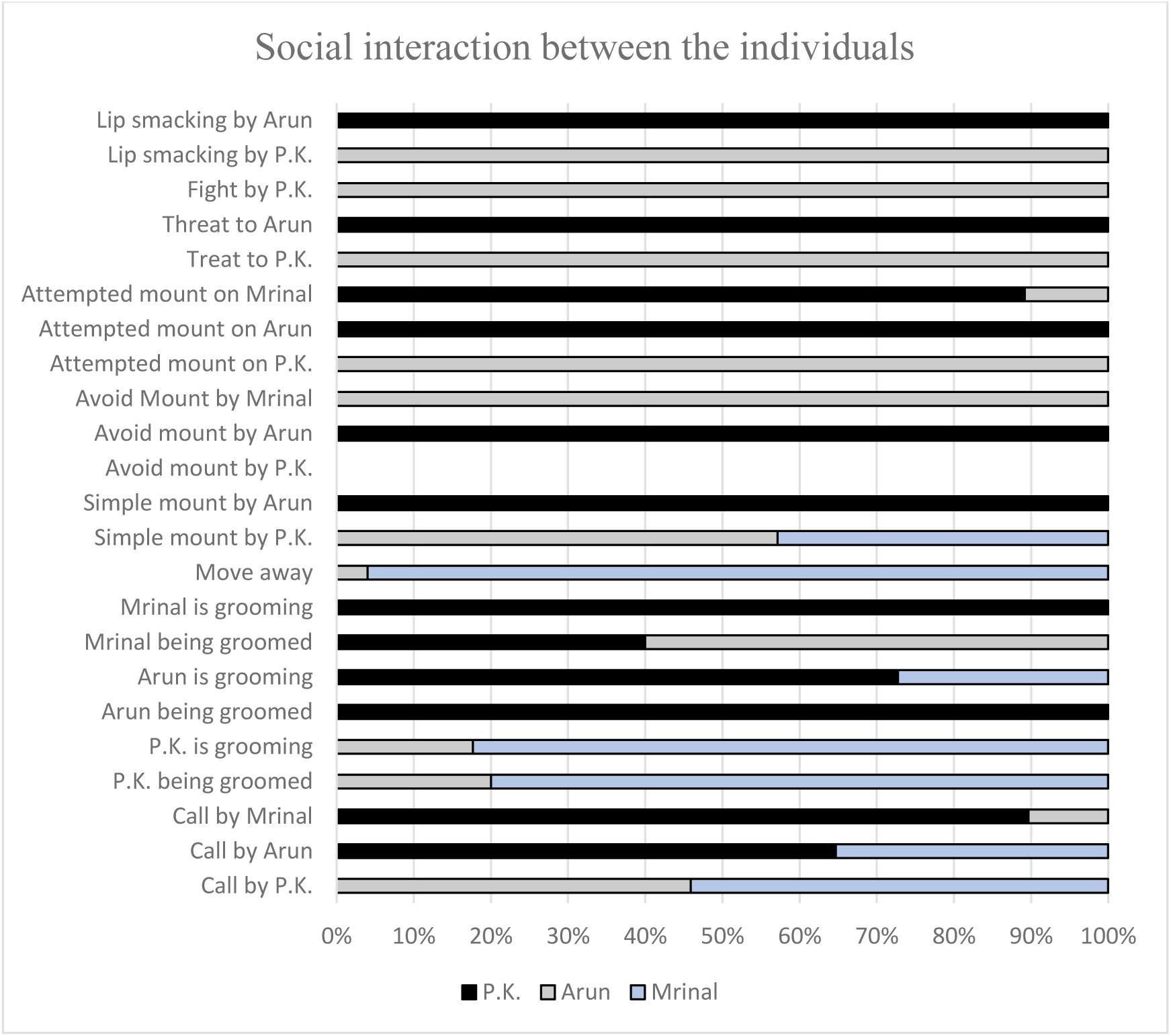
Social interaction between individuals living in NZP

## Discussion

Wild animals in captivity are often kept in conditions which are not suitable for their living and thus prevent them from exhibiting their natural species-specific behavioral patterns. Interestingly, it was seen that Arun depicted most of the abnormal trait who was confiscated from another zoo and was also deprived of any form of reproductive stimuli from the female. On the other hand, P.K. who was wild-caught exhibited few of the abnormal traits and displayed high rates of normal reproductive traits. Out of the abnormal behaviors, self-directed behaviors such as biting, masturbation, stereotype pacing, telescoping and rocking were exhibited only by the confiscated animal and almost never by the wild-caught individual. There are various studies which have been conducted on the developmental behavioral pathologies which have observed self-mutilating (e.g.- self-biting) and self-stimulating (e.g.- rubbing penis) behaviors to rise from an early stage in life (Anderson & Chamove, 1980, 1985; Chamove et al., 1984; Mallapur & Choudhury, 2003; Mallapur, Waran, et al., 2005a; Mootnick & Baker, 1994). It has been found that these forms of behavioral pathologies are a form of redirected social behaviors exhibited by an individual in the absence of social target (Chamove et al., 1984; Khan, 2013; King et al., 1980). These behaviors may be are the possible reason for the increase in the sensory inputs for the poor environmental conditions (Anderson & Chamove, 1985; Chamove et al., 1984; Mallapur, 2005b, 2005a; Mootnick & Baker, 1994).

A substantial degree of deviation was found when the ethogram of the captive lion-tailed macaques housed in NZP was compared with those in previous studies on the same species and in captivity (Skinner & Lockard, 1979). Although when compared with another study on captive lion-tailed macaques, a similar result was observed in the types of behaviors displayed by the captive animals in this study (Mallapur, 2005b; Mallapur & Choudhury, 2003; Mallapur, Waran, et al., 2005b, 2005a). Skinner and Lockard (1979) had not referred to any forms of abnormal behavior while discussing the behaviors displayed by the lion-tailed macaques in the captive environment during their study. This could either suggest that the abnormal behavior was not displayed by the animals or perhaps the observers had not recorded it. The major objective in the study conducted by Skinner and Lockard (1979) was to document the social interactions among individuals living in the group thereby putting more emphasis on social interactions and reproductive behavior rather than the abnormal behavior and welfare activities. In my study, I have tried to discuss the individual behavioral pattern both normal and abnormal behavioral traits displayed in captivity and discussed how behaviors varied with the animal’s rearing history and the social relations. The captive lion-tailed macaques observed in the Skinner and Lockard’s study (1979) were housed in a well-composed group whereas in the Indian zoos according to Mallapur (2005b) most of the animals were housed singly or in pairs. Thus, there is a high possibility that no abnormal behavior would be displayed by the animals in the former study. Most of the interactions on the social front (e.g. – lip smack, grooming, agonistic) were cited by Skinner and Lockard (1979) along with Mallapur’s study (2005b) on the captive lion-tailed macaques at the Indian zoos and few of them are also observed in this study. However, many features such as “head-bob”, “tail display”, “breast wipe” (Skinner and Lockard 1979), “regurgitate and re-ingest”, “oral stereotypy”, “eye flash”, “copulatory call” (Mallapur, 2005b; Mallapur, Waran, et al., 2005a) were not observed in this study. According to Skinner and Lockard (1979), the animal in the study had usually tossed their head at the ending of a threat display. They had hardly observed any behavior related to behavioral pathogens found by Mallapur (2005b).

When the study was compared using the ethograms for the free-ranging lion-tailed macaques few similar types of behaviors were observed (Hohmann, 1988; Raghavan, 2001). The lion-tailed macaques at the NZP were found to exhibit a number of behaviors such as auto-grooming, aggressive display, foraging which were also exhibited by free-ranging individuals (Raghavan, 2001). The studied individuals in this study were found to direct various social interactions such as affiliative teeth display, aggressive behavior and warning growls, which were reciprocated by or displayed to another individual. Although, similarity existed, on a broader view, the behavioral pattern of the lion-tailed macaque being studied were found to vary considerably from the free-ranging individuals and from other study conducted by other researchers in American zoo (Kumara et al., 2014; Raghavan, 2001; Singh, Kaumanns, Singh, Sushma, & Molur, 2009; Singh, Kumara, Kumar, & Sharma, 2001; Skinner & Lockard, 1979). As described earlier that the behaviors exhibited by animals living in captivity depend a lot on the artificial environments in which they are housed (Erwin, Maple, & Mitchell, 1979; Mallapur, Waran, et al., 2005a), the present study discusses the social environment i.e. social bond present between the members of the group. Captive primates required an appropriate social structure or group dynamics in order to endorse and exhibit their own species-specific behavior (Chamove et al., 1984; King et al., 1980; Rendall & Taylor, 1991). A study conducted by Mallapur (2005b) have found that factors such as the composition of the group, the enclosure design, the presence of visitors as well as the rearing history and the enrichment provided in the surrounding influence the behavior of the captive species.

Behavioral deprivation could also be influenced by most of the factors, such as group composition, the design of the enclosure and the feeding time. Primates, which are being housed singly or in inappropriate groups in the zoos results in a reduced normal or natural behaviors (Anderson & Chamove, 1980, 1985; Erwin et al., 1979; Rendall & Taylor, 1991). Studies on lion-tailed macaques living in their natural wild habitat have revealed that they live in multi-female (say 3 or 4) groups with infants and juveniles; usually 1 to 3 males (Kumar, 1987; Kumar et al., 2008; Raghavan, 2001). Housing captive individuals with the proper type of grouping system as present in wild results in decreased levels of stress among the individuals and incurring natural behavior (Skinner & Lockard, 1979). In another study conducted on captive animals displays the influence of group composition on their behavior which played a significant role in behavioral change (Kaumanns et al., 2013). Thus, the lacking of proper group dynamics in the study could have resulted in loose social interaction between the members as the female (Mrinal) highly preferred the alpha (P.K.). The result of weak social bonds resulted in the decreased reproductive success rate of the individual which could have resulted in increases in behavioral pathogens.

During this study, we have observed that the behavioral patterns of the captive lion-tailed macaques at NZP. In the study most of the behavioral pathogens have been observed in Arun who was confiscated, followed by Mrinal who was zoo-born and least in P.K. who was caught from the wild. The rearing history of the animals changed the lifestyle of the animals which was very clear from the types of behaviors displayed by them. Most of the behavioral abnormalities among the individuals at NZP have risen due to the absence of proper environmental stimuli such as species-specific group influence e.g. reproductive stimuli from Mrinal towards Arun. Deprivation from reproductive stimuli could have been the cause of the development of self-stimulatory behavioral pathologies. Thus, the NZP which have housed lion-tailed macaques as a part of long-term breeding programmes should strongly consider on their group dynamics as well as the rearing history of the animals while forming the group, so that the welfare of the animals is not compromised at any stage.

## Acknowledgements

We would like to thank the NZP authorities for granting the permission to conduct the research work. We would also like to thank Dr. Bhawal the veterinary officer of NZP for his constant support during the field work and Taniya Gill for her support during my fieldwork. The ethical permission to conduct this study was obtained from the University of Delhi and the proper permit was obtained from the NZP authorities. Our special gratitude goes to Department of Anthropology, the University of Delhi for providing me with funds to conduct the study.

